# Intercellular adhesion boots collective cell migration through elevated membrane tension

**DOI:** 10.1101/2024.07.03.601828

**Authors:** Brent M. Bijonowski, Jongkwon Park, Martin Bergert, Christina Teubert, Alba Diz-Muñoz, Milos Galic, Seraphine V. Wegner

## Abstract

In multicellular systems, the migration pattern of individual cells critically relies on the interactions with neighboring cells. Depending on the strength of these interactions, cells either move as a collective, as observed during morphogenesis and wound healing, or migrate individually, as it is the case for immune cells and fibroblasts. Transducers of cell-cell adhesions, such as cadherins coordinate collective dynamics by linking the cytoskeleton of neighboring cells. However, whether intercellular binding alone triggers signals that originate from within the plasma membrane itself, remains unclear. To address this question, we designed photoswitchable cell-cell adhesions that selectively connect adjacent plasma membranes without linking directly to cytoskeletal elements. We find that these intercellular adhesions are sufficient to achieve collective cell migration. Here, linking adjacent cells increases membrane tension, which activates the enzyme phospholipase D2. The resulting increase in phosphatidic acid, in turn, stimulates the mammalian target of rapamycin, a known actuator of collective cell migration. Collectively, these findings introduce a membrane-based signaling axis as promotor of collective cell dynamics, which is independent of the direct coupling of cell-cell adhesions to the cytoskeleton.

## Introduction

Collective cell migration is a hallmark of morphogenesis, tissue homeostasis, and wound healing ^1, 2, 3^. A central element that determines if cells move as a collective or not is the strength of intercellular adhesions ^4^. For instant, cell-cell adhesions mediated by classical cadherins link neighboring cells through the interactions of extracellular domains, while α-catenin and β-catenin recruit to the cytoplasmic tail to create a mechanosensitive link to the actin cytoskeleton ^5, 6^. Here, cadherins provide a mechanical link between cells that not only act as structural elements but also activate intercellular signaling circuits that alter cytoskeletal dynamics and the transcriptional state ^7, 8, 9, 10, 11, 12^.

In contrast to signaling circuits that emanate from protein complexes that link the cytoskeletons of adjacent cells to jointly determine the collective behavior, the contribution of signaling cues that originate from within the plasma membrane for collective cell dynamics has largely been overlooked. One reason for this is a lack of tools capable of dissecting the mechanical cues at the plasma membrane from the mechanotransduction to the cytoskeleton. For example, the truncation of the cytoplasmic catenin binding domains renders E-cadherin unfunctional, as the membrane-anchored extracellular domain by itself is insufficient to sustain intercellular cell-cell adhesions ^13, 14^. While this intricate mechanosensitive protein regulation at of cell-cell adhesions has precluded a detailed analysis in multicellular systems, the role of membrane properties for cell function has been well documented in single cells. Here, changes in membrane properties alter leading edge dynamics ^15, 16, 17^ and migration behavior ^18, 19, 20^. Noteworthy, alterations in membrane properties are accomplished not only through direct mechanical feedback loops, but also through intracellular signaling and gene expression ^21, 22^. As a result, changes at the plasma membrane not only yield short-term changes in cell dynamics but also have long-term effects ^23, 24^. While these findings suggest that membrane mechanics regulates cell dynamics of individual cells, the contribution of signals that originate in the plasma membrane and don’t require the direct link between adhesions and cytoskeleton in collective cell migration remains elusive.

Here, we set out to explore the relevance of membrane-based signaling for collective cell dynamics. To selectively elicit signals that originate from within the plasma membrane, we utilized artificial photoswitchable cell-cell adhesions based on the cyanobacterial phytochrome 1 (Cph1), which induce cell-cell adhesions on red light illumination without directly linking to the actin cytoskeleton ^25^. We found that an increase in cell-cell connections leads to coordinated and collective cell movement in cells that otherwise would migrate as single cells. Strikingly, increased cell-cell adhesions did not slow cells down, as would be expected when augmenting friction ^26^, but made them faster. At the cellular level, the photoactivated increase in cell-cell adhesions resulted in an increase in membrane tension. Importantly, it also activated the enzyme phospholipase D2, resulting in an increase in phosphatidic acid (PA) and activation of mammalian target of rapamycin (mTOR) signaling. These findings demonstrate that mechanical coupling of cells at their membranes via artificial adhesion proteins that do not link directly to the actin cytoskeleton induces increased membrane tension, which is sufficient to achieve collective cell migration.

## Results

### Cph1-PM-mediated transcellular adhesion alters collective cell migration

To explore the contribution of signals that originate at the plasma membrane in collective cell migration, we engineered artificial photoswitchable cell-cell adhesion molecules, which don’t link to the actin cytoskeleton. Specifically, we used as the extracellular domain of the Cph1 protein from *Synechocystis sp.*, which forms homodimers under red light (660 nm) and reversibly dissociates into monomers under far-red light (720 nm). To anchor Cph1 in the plasma membrane, without connecting to cytoskeletal components, Cph1 was solely fused to the transmembrane domain of the platelet-derived growth factor receptor (**Fig. 1a**). We named the membrane-anchored artificial and photoswitchable adhesion molecule ‘Cph1-PM’. In general, when Cph1- PM is expressed on the surfaces of cells, the adhesions between them will form under red light and reverse under far-red light as previously characterized ^25^.

**Figure 1.**
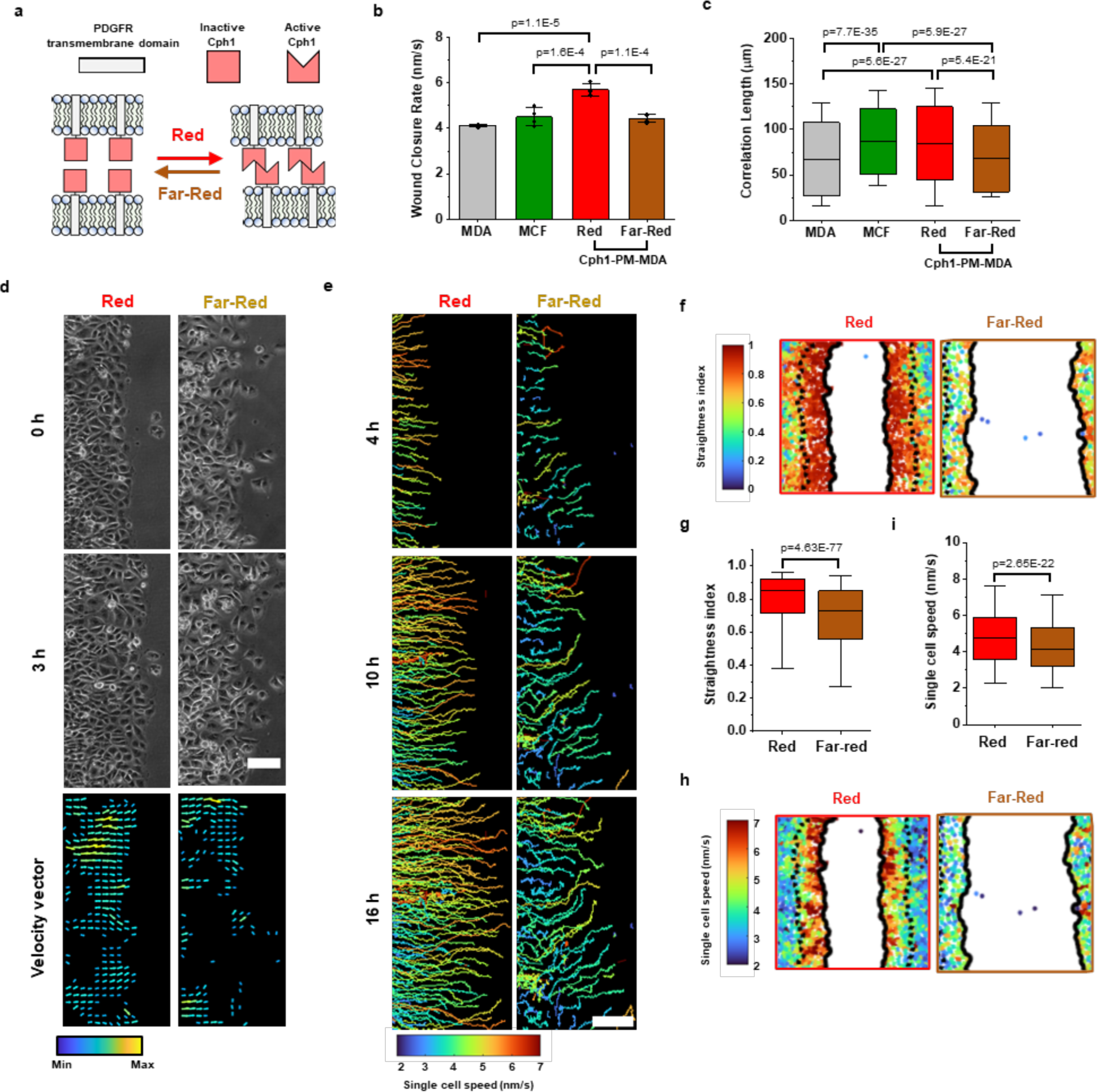
Cph1-PM activation alters cellular migration and coordination. (a) Schematic of Cph1-PM- based adhesion. **(b)** Average wound closure rate of parental MDA (n=4), MCF7 (n=4), Cph1-PM-MDA under red (n=4), and under far-red (n=4) from 4 biological replications (One-way ANOVA with Bonferroni’s multiple comparisons test). **(c)** Correlation length denotes the persistence distance of the velocity vector components of MDA (n=3443), MCF7 (n=793), Cph1-PM-MDA under red (n=732), and under far-red (n=1553) from 4 biological replications (One-way ANOVA with Bonferroni’s multiple comparisons test). **(d)** Phase-contrast micrographs (left) and vector velocity maps (right) depicting cell migratory trajectory of Cph1-PM-MDA cells under red (top) and far-red (bottom) light. **(e)** Single-cell tracking using nuclei to evaluate cell movement trajectory. Cell trace showing movement for first 5 hours. **(f)** Plots of Cph1-PM- MDA cell tracks with color code for straightness index under red or far-red light. **(g)** Straightness index, which evaluates the efficiency of wound healing, from 2 to 4 hours on Cph1-PM-MDA under red (n=2823) or far-red (n=2122) light from 2 biological replications (Two sample Mann-Whitney test). **(h)** Plots of Cph1- PM-MDA cell tracks with color code for single cell speed under red or far-red. **(i)** Single cell speed based on cell tracking from 2 to 4 hours under red (n=2377) and far-red light (n=1956) from 2 biological replications (Two sample Mann-Whitney test). Barplot (b) are denoted as the mean with standard deviation. Boxplots (c,g,i) present median, 25^th^ and 75^th^ percentile and the lower and upper boundaries of whiskers show 5^th^ and 95^th^ percentile. Scale bars, (d,e) 100 µm. Data source and statistic details are provided as a source data file.

As a starting point, we employed MDA-MB-231 (MDA) cells, which lack type-1 cadherins, display weak cell-cell adhesions and migrate as single cells ^27^. To explore if these artificial cell-cell adhesions can induce collective cell migration, we transfected MDA cells with Cph1-PM and generated monoclonal stable cell lines with Cph1-PM expression from them (Cph1-PM-MDA). In addition, we used MCF-7 cells as a positive control for a cell type that displays high expression levels of type-1 cadherins, strong cell-cell adhesions, and collective cell migration ^28^. To probe for collective and single-cell migration, we used wound healing assays and set a baseline for the migration behavior using wild-type MDA and MCF-7 cells. As expected, we find significantly faster the wound closure rates in MCF-7 cells compared to MDA cells (**Fig. 1b**). At the same time, MCF- 7 cells also display a higher correlation length than MDA cells, a measure of persistence distance and indicative of a higher degree of coordinated cell movement (**Fig. 1c**). We further observed individual MDA cells at the wound edge depart from the confluent cell sheet, indicative of the weak intercellular adhesions between them, while the leading edge of the MCF-7 wound remained intact (**Supplementary Fig. 1a** and **Supplementary Movie 1**).

Next, we probed the effect of increased intercellular adhesions on MDA cell migration. For this, we compared the cell migration behavior of Cph1-PM-MDA cells under red light, where the artificial cell-cell adhesions are active, to cells under far-red light, where the artificial cell-cell adhesions are inactive. Exposure to red light caused Cph1-PM- MDA cells to migrate collectively and with a unified front (**Fig. 1d** and **Supplementary Movie 2**). Directional rate vectors for the migration showed that under red light the majority of motion vectors were directed toward the wound space. On the other hand, the orientation of the rate vectors of Cph1-PM-MDA cells under far-red light and wild-type MDA cells were random (**Supplementary Fig. 1a**). Moreover, we found a significantly higher correlation length for Cph1-PM-MDA cells under red light than under far-red light, which had a comparable correlation length to the parent MDA cells (**Fig. 1c**). Actually, the correlation length pf Cph1-PM-MDA cells under red light and MCF-7 cells were comparable.

Counterintuitively, the wound closed faster for Cph1-PM-MDA cells under red light than either those under far-red light or MDA cells lacking the construct (**Fig. 1b**), despite increased friction between the cells and the cells being more restricted in their freedom to move independently ^26^. These differences in wound closure rate were consistent over the entire 12-hour duration of the experiment (**Supplementary Fig. 1b**). To further analyze the origin of the faster wound closure rate under red light, we performed single-cell motion analysis (**Fig. 1e** and **Supplementary Movie 3**). The traces of individual Cph1-PM-MDA cells under red light exhibited a directional migration towards the wound during the analyzed 16-hour time window, whereas Cph1-PM-MDA cells under far-red light showed frequent random trajectories. For analysis, cells were grouped based on localization into “front cells”, (i.e. 3 cell diameters from the wound edge) and “rear cells” (**Supplementary Fig. 2a**). Under red light, Cph1-PM-MDA cells at the “front” moved more directly towards the wound compared cells under far-red light (**Figs. 1f, g, Supplementary Figs. 2b, c**). Moreover, also at the single cell level Cph1-PM-MDA cells at the front migrated faster under red light than under far-red light (**Fig. 1h**).

### Cph1-PM mediated transcellular adhesion augments plasma membrane tension

Increasing the friction between individual cells was previously reported to slow down the median speed while increasing the collectivity of individual agents ^26^. In stark contrast, we found that increased intercellular adhesion in Cph1-PM-MDA cells augments cell speed and directionality at the same time (**Fig. 1b, h**). These findings suggest that Cph1-PM activation does not merely increase the adhesion between adjacent cells, but that it also elicits downstream cell signaling. Since Cph1-PM lacks a cytosolic tail that could directly interact with intracellular proteins, we hypothesized that this signaling must originate from within the plasma membrane. To test this assumption, we probed for changes in membrane properties.

First, we measured whether Cph1-PM activation influenced lipid ordering. For that, we took advantage of the hydrophobic fluorescence membrane dye Pro12A, which exhibits a blue-shifted emission with increased lipid ordering. In this ratiometric imaging, a rise in generalized polarization (GP) is associated with stiffer gel-like lipid conditions and a decrease is associated with increased fluidity ^29^. Focusing on a single cell-cell interface, we observed an increase in GP when activating the Cph1-PM with red light for 15 min (**Fig. 2a**). Opposingly, when the cells were exposed first to red light for 15 min and then for 15 min to far-red light illumination, to inactivate the Cph1-PM based adhesions, the GP value decreased (**Fig. 2b**). The results indicated that the direct physical cell-cell membrane adhesions locally alter membrane structure and make the interphase membranes stiffer at the cell-cell contact sites.

**Figure 2.**
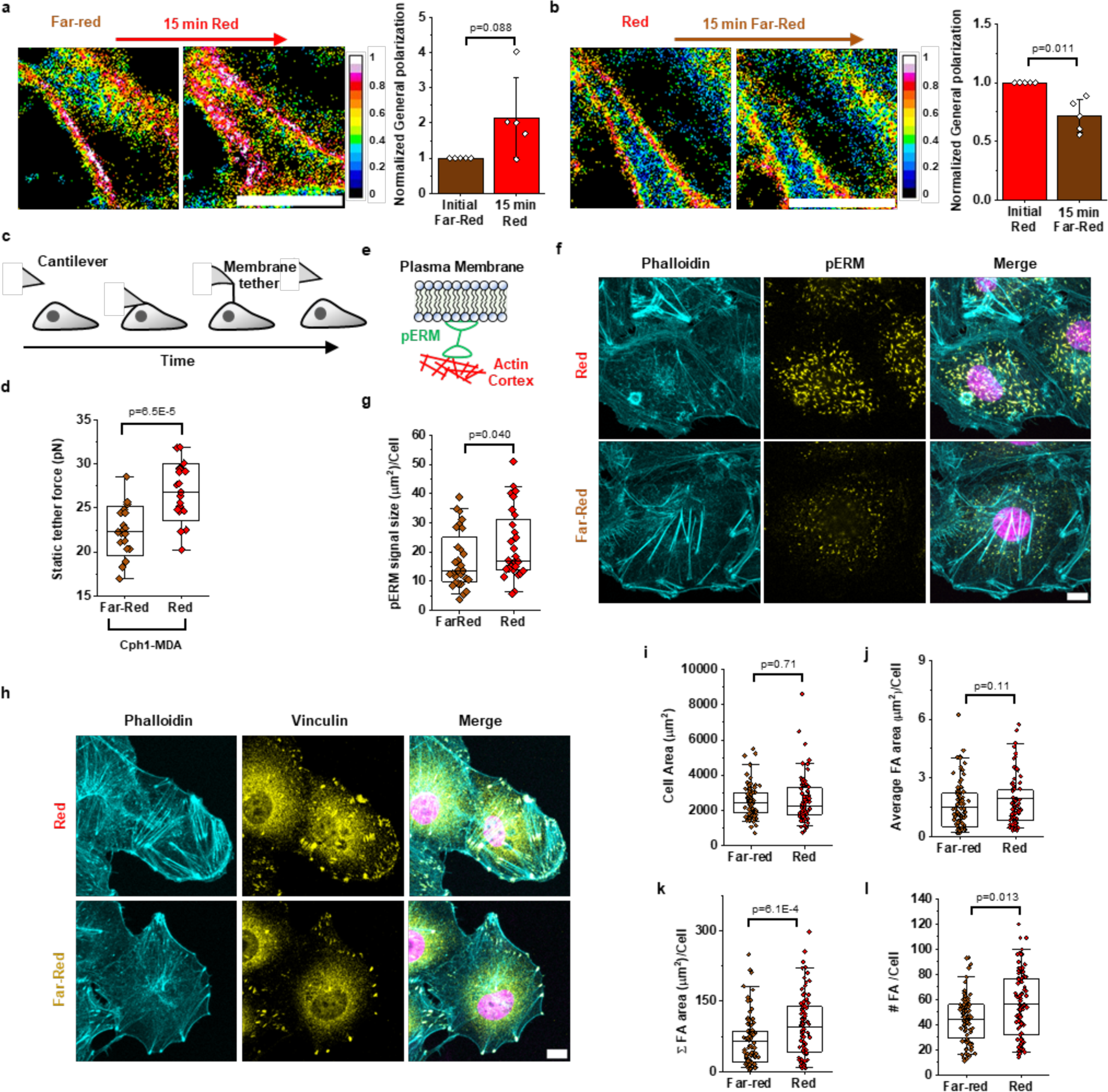
Activation of Cph1-PM augments membrane tension, ERM, and focal adhesion expression. (a, b) Pro12A staining shows polarization changes upon changing illumination from far-red to red (a) and from red to far-red (b) light. Plots depicting the normalized general polarization at the cell-cell interface upon changes from far-red to red light (a; n=5) or from red to far-red (b; n=5). (n=2 biological replications; Pair sample t-test). **(c)** Schematic showing how the static tether pulling force is measured by AFM. **(d)** Static tether force of Cph1-PM-MDA cells exposed to far-red (n=20) or red (n=20) from 3 biological replicates. **(e)** Schematic of pERM proteins. **(f)** Immunostaining of Cph1-PM-MDA cells stained with phalloidin (cyan), pERM (yellow), and DAPI (violet). **(g)** Quantification of pERM signaling per cells upon exposure to far-red (n=32) and red (n=32) from 3 biological replications (Two samples Mann-Whitney test). **(h)** Immunostaining of Cph1-PM-MDA cells stained with phalloidin (cyan), Vinculin (yellow), and DAPI (violet). **(i-l)** Cell spreading area (j), average FA area per cell (k), total FA area per cell (l), and FA number per cell (m) of Cph1-PM-MDA cells exposed to either far-red (n=67) or red (n=69) from 3 biological replications (Cell area: two sample t-test, average FA area per cell and total FA area per cell: two sample Mann-Whitney test, FA number per cell: two sample t-test). Barplots (a,b) are denoted as the mean with standard deviation. Boxplots (d,g,I,j,k,l) present median, 25^th^ and 75^th^ percentile and the lower and upper boundaries of whiskers show 5^th^ and 95^th^ percentile. Scale bars, (a,b,f,h) 10 µm. Data source and statistic details are provided as a source data file.

To further analyze the alternations in membrane properties on induction of the artificial cell-cell adhesions, we performed static tether pulling using atomic force microscopy (AFM) to examine changes in apparent plasma membrane tension at the cellular scale. Specifically, a membrane tether on cells positioned close to the wound edge was maintained at a constant distance until it ruptures, and the corresponding force difference was quantified (**Fig. 2c**). We found that Cph1-PM-MDA cells had a significantly higher static tether force under red light than these cells under far-red light (**Fig. 2d**). Moreover, arguing against profound tension gradients between different cells, we did not observe significant differences in membrane tension in the front vs. rear of the wound for either condition (**Supplementary Figs. 2a, b**).

### Cph1-PM mediated changes in membrane tension affect membrane-to-cortex attachment and formation of focal adhesions

The static tether force measured in the previous experiments is proportional to the apparent membrane tension, which integrates the in-plane tension of the lipid bilayer as well as the membrane-to-cortex attachment ^19, 30, 31^. To investigate the origin of the observed difference in apparent membrane tension, we next probed the membrane-cortex interface. Previous studies showed that phosphorylation of Ezrin/Radixin/Moesin proteins (pERM) increases the connection between F-actin and the plasma membrane, thereby increasing the apparent membrane tension (**Fig. 2e**) ^32^. To probe for changes in pERM phosphorylation upon Cph1-PM activation, we exposed Cph1-PM-MDA cells to either red or far-red light, fixed the cells under the respective illumination, and then stained for F-actin, pERM, and DAPI (**Fig. 2f**). Here, we find quantitative increase of the pERM signal in Cph1-PM-MDA cells under red compared to far-red light (**Fig. 2g**) as another indicator of increased membrane tension on formation of the artificial cell-cell adhesions.

Considering that membrane tension has been reported to impact focal adhesion in single cells ^33^, we investigated how cell membrane adhesion affected cell spreading and the number of focal adhesions ^34, 35^. We found an increased number of focal adhesions in Cph1-PM-MDA cells under red light compared to those under far-red light by staining the focal adhesion associated protein vinculin, which is only associated with focal adhesions in the Cph1-PM-MDA system (**Figs. 2h, l**). While the total area of the focal adhesions per cell increased, the cell spreading area and the area of individual focal adhesions between Cph1-PM-MDA cells under red and far-red light did not show significant differences (**Figs. 2i, j, k**). Previous reports showing that not the total number but the size of the focal adhesions are the determining factor of cell speed would argue that the increased number of cell adhesions under red light compared to far-red light would not slow down cell migration ^36^.

### Cph1-PM mediated effects are independent of E-cadherin signaling

Having observed collective migration, a feature generally associated with epithelial cell types, in the Cph1-PM-MDA originating from a mesenchymal cell type (MDA), we investigated changes in the expression of epithelial and mesenchymal markers. First of all, we found no increase in the expression of the cell-cell adhesion molecule E-cadherin (marker of the epithelial phenotype) in Cph1-PM-MDA cells independent of light treatment, while it was clearly expressed in control MCF7 cells (**Fig. 3a**).

**Figure 3.**
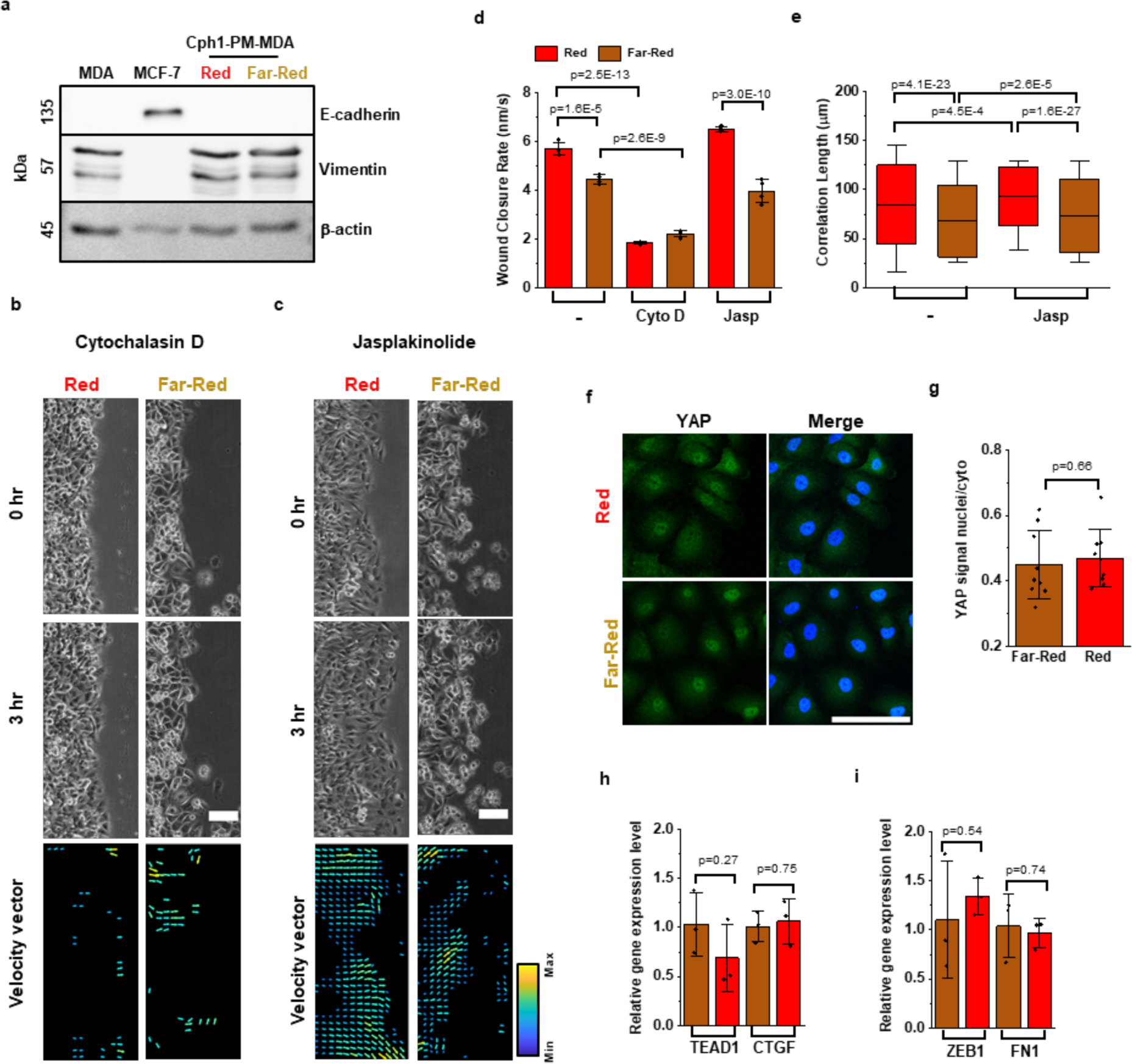
Cph1-PM-dependent changes in collective cell dynamics do not change cell identity. (a) Western Blot analysis of E-Cadherin and Vimetin in parental MDA cells, MCF cells, as well as in Cph1-PM- MDA cells exposed to red and far-red light. **(b, c)** Phase-contrast micrographs and vector velocity maps for Cph1-PM-MDA cells under red and far-red light upon the addition of CytoD (a) and Jasp (b). **(d)** Average wound closure rate of Cph1-PM-MDA without treatment under red (n=4) and far-red (n=4), with Cyto D under red (n=4) and far-red (n=4), or with Jasp under red (n=4) and far-red (n=4) (One-way ANOVA with Bonferroni’s multiple comparisons test). **(e)** Correlation length denotes the persistence length of the velocity vector of Cph1-PM-MDA under red (n=732) or far-red (n=1553) light, with Jasp under red (n=533) or far-red (n=2440) (n= 4 biological replications; One-way ANOVA with Bonferroni’s multiple comparisons test). **(f)** Immunofluorescent images of Cph1-PM-MDA cells under red (top) and far-red (bottom) light stained against YAP (green) and DAPI (blue). **(g)** Quantification of YAP localization in Cph1-PM-MDA cells under far-red (n=9) and red (n=9). Plot depicts the nuclear vs cytoplasmintensity. (3 biological replications, two sample t-test). **(h)** Gene expression of TEAD1 and CTGF in Cph1-PM-MDA cells exposed to far-red (n=3) and red (n=3) light (n= 3 biological replications, two sample t-test). **(i)** Gene expression levels of ZEB1 and FN1 in Cph1-PM-MDA exposed to far-red (n=3) and red (n=3) light (Two sample t-test). Barplots (d,g,h,i) are denoted as the mean with standard deviation. Boxplots (e) present median, 25^th^ and 75^th^ percentile and the lower and upper boundaries of whiskers show 5^th^ and 95^th^ percentile. Scale bars (b,c), 100 µm; (f), 20 µm. Data source and statistic details are provided as a source data file.

The observed collective migration in Cph1-PM-MDA cells under red light and the faster movement of individual cells raises the question of what signaling pathways are triggered downstream. A logical starting point was to investigate whether signaling pathways that are associated with cadherin signaling are indirectly activated due to these artificial cell-cell adhesions. First, to determine the role of actin polymerization in Cph1-PM dependent signal propagation, we added cytochalasin D (CytoD), an inhibitor of actin polymerization (**Fig. 3b**) and jasplakinolide (Jasp), an actin nucleation promoter (**Fig. 3c**) to the wound healing assays. Treatment with CytoD reduced the wound closure rate both in Cph1-PM-MDA cells under red and far-red light, only confirming the central role of actin in cell migration (**Figs. 3b, d**). Moreover, the addition of Jasp yielded no significant changes compared to the untreated cells (**Figs. 3c, d**), except for a slight increased correlation length for Cph1-PM-MDA cells both under red and far-red light when treated with Jasp (**Fig. 3e**).

Next, we evaluated Cph1-PM-dependent changes in the mechanosignaling associated with E-cadherin. E-Cadherin has been described to activate YAP-dependent gene expression with changes in mechanical strain ^37^. When evaluating the localization of YAP under different illumination, we found no difference in the cytoplasmic versus nuclear localization (**Figs. 3f, g**). Consistently, expressions of two YAP down-streaming markers, TEA domain transcription factor 1 (TEAD1) and connective tissue growth factor (CTGF), showed no significant differences between red and far-red (**Fig. 3h**).

Finally, we evaluated EMT markers that would be expected to be down regulated with increasing epithelial characteristic. Firstly, vimentin, a mesenchymal cell marker, showed no change in the Cph1-PM-MDA compared to the parent cell type in the western blot (**Fig. 3a**). Likewise, the gene expression levels of two representative mesenchymal markers, zinc finger E-box binding homeobox transcription factor 1 (ZEB1) and fibronectin 1 (FN1), were comparable with red and far-red light exposure as observed with RT-PCR (**Fig. 3i**). Collectively, we did not find any evidence that increased Cph1-PM based cell-cell adhesions with red light exposure alters the expression of genes that are associated with the transition from a mesenchymal to an epithelial cell type. Hence, these findings argue that Cph1-PM signaling and the observed collectivity in the wound healing did not rely on any cadherin-dependent circuits.

### Cph1-PM mediated transcellular adhesion promotes PLD2 signaling

The findings to this point establish that activation of the artificial cell-cell adhesions results in substantial changes in membrane tension as well as in faster and collective cell migration. At the same time, we find no changes in signaling cascades originating from classical E-cadherin signaling. Unlike E-cadherin with truncated cytosolic tail, which cannot mediate cell-cell adhesions due to the missing interactions with the cytoskeleton, Cph1-PM is able to maintain stable cell-cell adhesions under red light without connecting to the cytoskeleton. Therefore, we reasoned that the origin of the signal resulting in the altered phenotype may reside in the plasma membrane. Indeed, membrane tension has been implicated promoting biochemical signals at the single-cell level ^20^. More precisely, increased membrane tension was reported to augment phospholipase D2 (PLD2) activity, which breaks down phosphatidylcholine (PC) into phosphatidic acid (PA) and choline ^20^. Downstream, PA modulates the properties of cells through activation of various pathways, chief among others, is the mammalian target of rapamycin (mTOR) pathway ^38^. Considering its mechanosensitive properties, we thus probed for changes in PLD2 activity, through the quantification of its product PA ^21^. Consistent with increased PLD activity, we observed significantly elevated PA levels in Cph1-PM-MDA cells under red light compared to cells under far-red light and the MDA cells lacking Cph1-PM (**Fig. 4a**). This increase in PA in Cph1- PM-MDA cells under red light was clearly due to the increased cell-cell adhesions and not the expression of Cph1-PM in MDA cells itself as no significant differences in PA content was observed between Cph1-PM-MDA cells under far-red light and the parent MDA cells.

**Figure 4.**
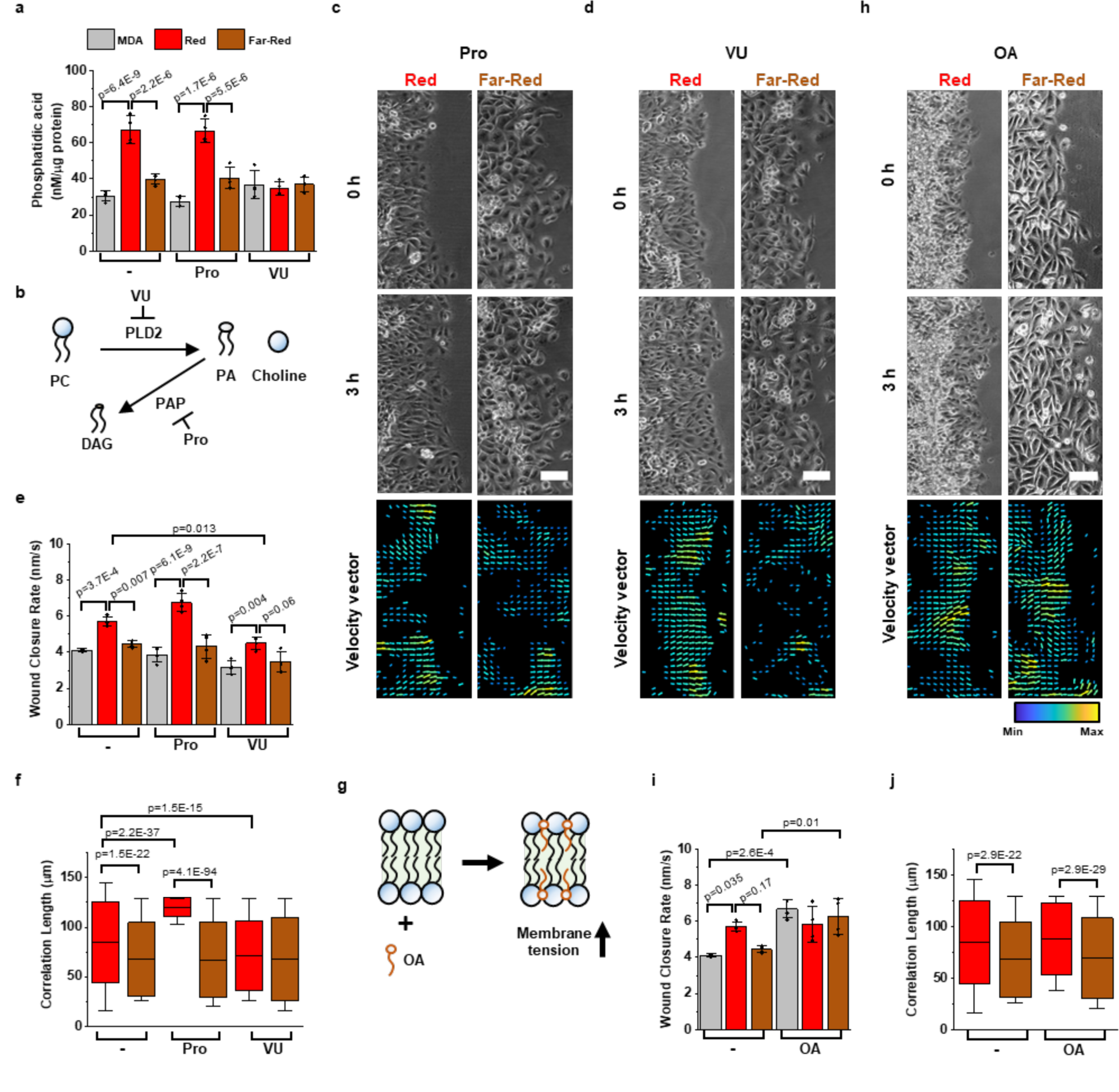
Cph1-PM-dependent changes in collective cell dynamics are mediated by PLD2 activity and membrane tension. (a) Phosphatidic acid levels for parental MDA (gray graphs) and Cph1-PM-MDA cells under red (red graph) and far-red (brown graph) in the absence or upon addition of Pro or VU (All data n=4 from 4 biological replications) **(b)** PLD2 hydrolyzes phosphadic choline into phosphatidic acid (PA) and choline and phosphatidic acid phosphatase (PAP) hydrolyzed PA to diacylglycerol (DAG). PAP is inhibited by Pro and PLD2 is inhibited by VU. **(c, d)** Representative phase-contrast micrographs and histograms of vector directionality for Cph1-PM-MDA cells under red and far-red light after treatment of Pro (c) and VU (d). **(e)** Wound closure rate from parental MDA or Cph1-PM-MDA cells under red and far-red light. For each condition, cells are shown before treatment or after treatment with Pro or VU. (All data n=4 from 4 biological replications) **(f)** Correlation length of Cph1-PM-MDA cells without treatment under red (n=732) and far-red (n=1553) light, or after treatment of Pro under red (n=252) and far-red (n=1588) light or after treatment of VU under red (n=2213) and far-red (n=1209) light from 4 biological replications. **(g)** OA incorporated into the lipid bilayer of cell membranes and increases the membrane tension. **(h)** Phase-contrast micrographs and velocity vector map for Cph1-PM-MDA cells under red and far-red light after treatment of OA. **(i)** Wound closure rate from parental MDA cells or Cph1-PM-MDA under red and far-red light without treatment or after treatment with OA. (All data n=4 biological replications) **(j)** Correlation length denotes the persistence length of velocity vector under red and far-red before and after treatment with OA (no treatment, n_red_=732, n_far-red_=1553; OA, n_red_=846, n_far-red_ =1581; 4 biological replications). All statistic was analyzed by one-way ANOVA with Bonferroni’s multiple comparisons test. Barplots (a,e,i) are denoted as the mean with standard deviation. Boxplots (f,j) present median, 25^th^ and 75^th^ percentile and the lower and upper boundaries of whiskers show 5^th^ and 95^th^ percentile. Scale bars (c,d,h), 100 µm. Data source and statistic details are provided as a source data file.

To further consolidate these findings, we modulated the PLD2 pathway by adding either propranolol (Pro), which prevents the degradation of PA through phosphatidic acid phosphatase (PAP)^39^, or by adding VU-0285655-1 (VU), which inhibits PLD2 directly (**Fig. 4b**) ^40^. Following treatment with Pro, PA levels in Cph1-PM-MDA cells under red light remained as high as for untreated cells (**Fig. 4a**). In contrast, the addition of VU decreased PA levels in Cph1-PM-MDA cells exposed to red light to the level of cells under far-red light. Other than this, no significant changes in PA levels were observed when Cph1-PM-MDA cells were treated with either Pro or VU under far-red light or in parent MDA cells. These findings are relevant, as it argues that no additional PA is produced in Cph1-PM-MDA cells under red light illumination without the activation of PLD2 (**Fig. 4a**).

To connect the increases in PA with the increased migratory capacity as has been demonstrated for single cells, ^20, 31, 41^ we evaluated the effect of altered PLD2 activity on wound closure rates and coordination of the cellular movement (**Figs. 4c, d**). In the presence of Pro, Cph1-PM-MDA cells under red light showed similar wound closure rates as Cph1-PM-MDA cells under far-red light and untreated MDA control cells, respectively. In contrast, the treatment with VU resulted in a significant decrease in wound closure rate under red light, while it did not alter the wound closure rate under far-red light (**Fig. 4e**). This result directly indicates that PLD2 activity and PA levels contribute to the improved wound closure rate and faster movement of individual cells observed for Cph1-PM-MDA cells under red light. This relationship is further supported by the decreased correlation length with VU treatment under red, which comparable to the correlation length of Cph1-PM-MDA cells under far-red light (**Fig. 4f**). Collectively, these results suggest not only that the PLD2 activation modulates the migration potential of MDA cells, but also that Cph1- PM activates the PLD2 pathway.

Finally, we increased the membrane tension in cells through the addition of oleic acid (OA), which inserts itself into the plasma membrane resulting in a uniform increase in membrane tension (**Fig. 4g**). Exogenous increase of membrane tension with OA led to Cph1-PM-MDA cells under far-red light and parent MDA cells to migrate at the same pace as Cph1-PM-MDA cells under red light (**Figs. 4h, i**). However, it should be noted that under far-red light in the presence of OA, the Cph1-PM-MDA cell migration was not as coordinated as for the cells under red light due to the lack of cell-cell connections (**Fig. 4j, Supplementary Video 4**).

### Cph1-PM mediated membrane signaling impacts the PI3K/mTOR pathway

The results to this point suggest that the direct connection between cells through Cph1-PM increased membrane tension, activating PLD2, which in turn increased PA levels. To understand how this may affect cell dynamics, we investigated the role of PA-associated pathways. In particular, we considered the PA-dependent PI3K/mTOR activity shown to impact cell migration^42^. To this end, we perturbed the PI3K/mTOR pathway with wortmannin, an inhibitor of PI3K ^43^; SF1670, an inhibitor of PTEN ^44^; and rapamycin, an inhibitor of the mTOR complex ^45^. After wortmannin treatment, the wound closure rate of Cph1-PM-MDA cells under red light significantly decreased and was comparable to cells under far-red light (**Fig. 5a**). Similarly, the addition of rapamycin led to a significant decrease in the wound closure rate of Cph1-PM-MDA cells under red light, but had not affect under far-red light. In contrast, SF1670 treatment resulted in a significant increase in the wound healing rate even under far-red light, but did not alter Cph1-PM- MDA cells under red light (**Fig. 5b**). Complementarily, we evaluated single-cell tracks during the wound healing assay under red and far-red light in conjunction with SF1670 and rapamycin treatment (**Fig 5c, Supplementary Figs 4a, b**). In the absence of any inhibitor, as already observed, the cell trajectories were straighter and the individual cells moved faster under red light than under far-red light. Yet, the SF1670 treatment resulted in an increased coordination of the cells even under far-red light. These findings are consistent with previous studies that reported an increase in the wound healing rate, and a tendency for collective cell movement, upon inhibition of PTEN ^46^. Importantly, the wound closure rate decreased for Cph1-PM-MDA cells treated with rapamycin under red, but not under far-red light, which argues that inhibition of the mTOR pathway selectively suppressed the collective behavior of the Cph1-PM-MDA cells induced by the artificial cell-cell adhesions.

**Figure 5.**
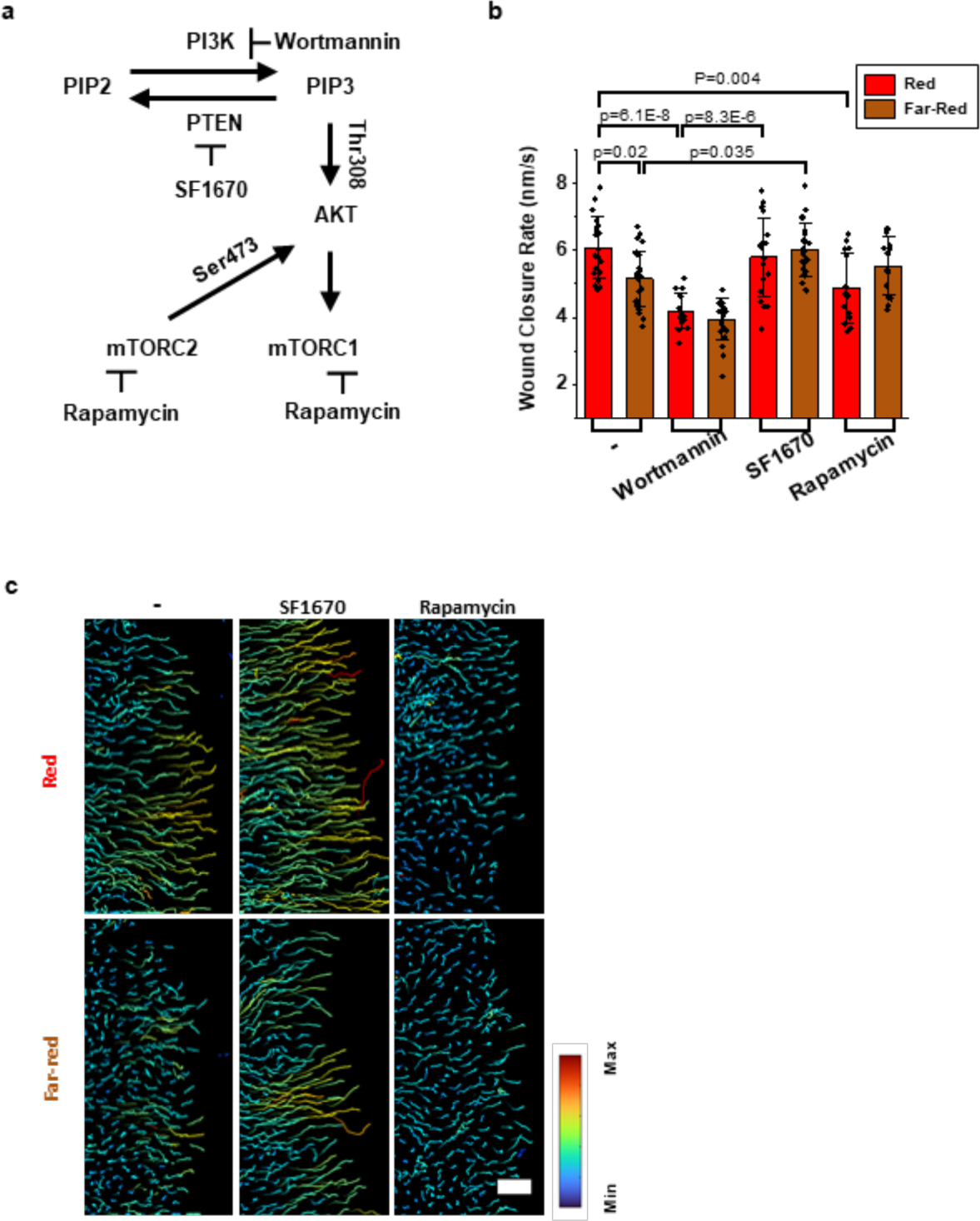
Cph1-PM-dependent changes in collective cell dynamics are mediated through PI3K/mTOR signaling pathway. (a) Schematics of PI3K and mTOR signaling pathway. PI3K is inhibited by Wortmannin, PTEN is inhibited by SF1670, and mTORC1 and mTORC2 are both inhibited by rapamycin. **(b)** Migration rate over time and the average rate under red and far-red illumination (no treatment, n_red_=20, n_far-red_=24; Wortmannin, n_red_ =16, n_far-red_ =23; SF1670, n_red_=19, n_far-red_ =22; Rapamycin, n_red_ =14, n_far-red_=10; from 3 biological replications; One-way ANOVA with Bonferroni’s multiple comparisons tests). **(c)** Single- cell tracking using nuclei showing cell trajectories over 5 hours. Scale bars, 100 µm. Bar plots (b) are denoted as the mean with standard deviation. Data source and statistic details are provided as a source data file.

## Discussion

Intercellular coupling is crucial for the emergence of collective cell dynamics. It therefore does not surprise that several types of cell-cell adhesion proteins facilitate this intercellular cohesion. For instance, cadherins at adherent junctions and claudins/occludins at tight junctions bridge the actin cytoskeleton of adjacent cells, while desmosomal proteins link intermediate filaments between cells ^47^. Furthermore, although primarily known for cell-extracellular matrix interactions, integrins were recently shown to also influence cell-cell adhesions ^48^. While distinct in structure, localization and function, many of these proteins display not only adhesive properties but also mechanosensitive cell signaling. For instance, stretching of alpha-catenin, which links adherent junctions to actin filaments, exposes a cryptic binding sites for vinculin ^49^. Similarly, talin, which participates in transferring ECM-based forces at focal adhesions to actin filaments, exposes under load a binding site for vinculin ^50^. Intriguingly, recent studies showed that desmoplakin, which couples desmosomes to intermediate filaments, also undergoes conformational changes under mechanical stress ^51^. The picture emerging from these and other studies is that differences in the expression level of these proteins define the distinct cell-specific adhesive and mechanosensitive signaling properties, which jointly determine the collective cell dynamics.

Complementing the current protein-centric view, we explored in this study the role of the plasma membrane in this process. By using the artificial photoswitchable Cph1-PM-based cell-cell connections, we ensured that the differences in migration only arise from the connections between the cells and not from some other genetic or environmental factors. We find that augmented cell-cell adhesion mediates membrane ordering and a stiffer interface. The resulting increase in membrane tension promotes signaling. Specifically, we demonstrate that activation of PLD2 augments PA cleavage, which activates the mTOR/PI3K pathway to promote cell dynamics. These findings are consistent with experimental work obtained in single cells ^20^. Hence, frictional forces at the plasma membranes initiate, analogous to mechanosensitive proteins under mechanical stress, a signaling cascade to synchronize collective cell dynamics (**Fig. 6**).

**Figure 6.**
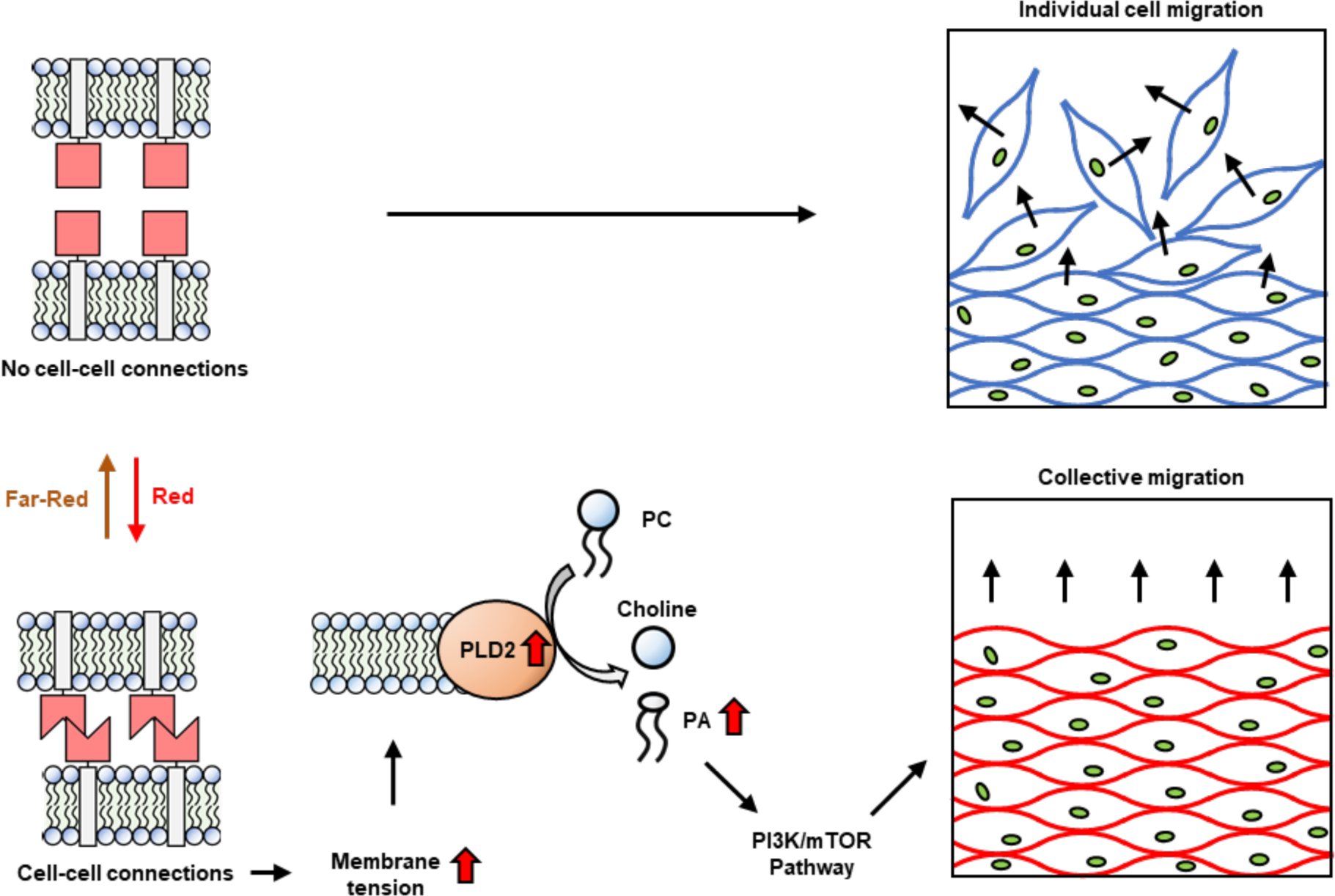
Schematic of the proposed mechanism. When cells are not connected, they migrate randomly. As cells become adherent, the physical adhesion increases cell membrane tension. The increase in membrane tension, in turn, promotes PLD2 activity, resulting in the degradation of PC into PA. The elevated PA activates the PI3K/mTOR pathway, promoting collective migration.

Collectively, these findings establish that cell-cell connections alter membrane mechanics, which in turn alters collective migration. We consider this another regulatory layer that acts in parallel to protein-based systems to coordinate collective cell dynamics. Given that Cph1-PM can be functionally replaced by any transmembrane protein, it is plausible to envision that the identified signaling circuit is of relevance for a broad spectrum of phenomena where mechanical stress is exerted on the plasma membrane.

## Methods

### Construct and sequence

The Cph1 domain of Cph1-PM was cloned into a pDisplay mammalian expression vector (Invitrogen V66020) between the Ig κ-chain leader sequence and the platelet-derived growth factor receptor domain as a Cph1-GFP fusion as previously described by our group ^25^. The N-terminal to the murine Ig κ-chain leader sequence directs the protein to the secretory pathway, while the C-terminal PDGFR transmembrane domain anchors the proteins on the extracellular side of the plasma membrane.

### Cell culture

MDA-MB-231 (HTB-26) and MCF-7 (HTB-22) cells were both purchased from ATCC. All cells were cultured in DMEM (Dulbecco’s Modified Eagle Medium, Gibco) supplemented with 10% FBS (fetal bovine serum, Gibco), 1% penicillin/ streptomycin (Gibco), and 12.5 mM HEPES (Sigma Aldrich) at 37 °C and 5% CO_2_. Cells were subcultured when they reached 80% confluence. A monoclonal stable cell line, Cph1-PM-MDA, that expresses Cph1-PM was generated starting from MDA-MB-231 cells as previously described ^25^. Cph1-PM-MDA cells were maintained in 1800 μg/mL G418 (Geneticin, Roche) and used for up to 10 passages.

### Wound healing assay

The wound-healing assay was performed in 24-well plates (Greiner Bio-One), into which MCF-7, MDA-MB-231, or Cph1-PM-MDA cells were seeded at 1.5x10^5^ cells per well. Phycocyanobilin (PCB, 5 μM) was added to the Cph1-PM-MDA cells, and incubated overnight at 37 °C and 5% CO_2_. Note that the activation of Cph1-PM is dependent on the presence of its cofactor PCB, as the protein remains inactive under red light in the absence of PCB treatment. 4 hours before imaging, a vertical wound was created with a 200 μL tip, and fresh media containing the drugs (DMSO (control), Cytochalasin D (0.1 μM), Jasplakinolide (25 nM), Propranolol (1 μM), VU- 0285655-1 (5 μM), oleic acid (50 μM), wortmannin (0.2 μM), SF1670 (2 μM), or rapamycin (0.2 μM) was added. No additional PCB was added, since it would aggregate with the oleic acid and Cph1-PM domains were already saturated.

### Microscopy

An hour before imaging the multiwall plate was moved to an Ibidi multiplate heating system with gas control. The system was configured according to the manufacturer specifications (bottom glass 37 °C, cassette 38 °C, and top glass 42 °C), and maintained at 5% CO_2_. Images were taken on an inverted Leica DMi8 microscope, using a 10x phase-contrast objective, every 10 minutes for 16 hours. A 533/SP Bright Line HC short-pass filter (AHF Analysentechnik, 380-520 nm transmission) was used in front of the white light source to avoid photoactivation. Illumination was achieved with either an external red or far-red light lamp, which provided continuous illumination The light intensities were 1440 μW/cm^2^ with 620 nm for red light and 1120 μW/cm^2^ with 734 nm for far-red light.

### Image Analysis

For wound healing, images were analyzed with ImageJ (National Institute of Health, USA) to determine the area of the wound at each time point. The zero-time point was set at 4 hours post wounding to avoid the effects resulting from variation in the cell seeding density and wounding process. Further vector analysis was carried out in MATLAB version 7.10 (R2020a; Mathworks, USA) and Particle image velocimetry (PIV) plugin from Dr. William Thielicke and Prof. Eize J. Stamhuis (MATLAB plugin). Custom code was used to create colored vector plots and correlation lengths ^9^.

For single-cell tracking analysis, Cph1-PM-MDA cells were stained with 0.5 μM Hoechst 34580 for 15 minutes and a wound was created with a 200 μL tip. Following, the cells were washed twice with PBS, and the media was replaced with DMEM containing 10% FBS with/without drugs. The Ibidi multiplate heating system and DMi8 microscope with the same settings as above (see “Microscope”) were employed for image acquisition. The active form is of Cph1-PM is measured in the presence of PCB under red light, while the inactive form measured is in the presence of PCB under far-red light or without PCB under red light. Bright-field and nucleus images at a wavelength of 405 nm were captured every 10 minutes over a period of 16 hours.

### Pro12A staining

Pro12A was generously gifted by Dr. Andrey S. Klymchenko, University of Strasbourg. Cph1-PM- MDA cells were seeded at 1x10^4^ cells/cm^2^ into a glass coverslip bottomed Ibidi m-dish 35 mm with PCB cofactor and incubated overnight to ensure adhesion. 45 minutes before imaging, 5 μM Pro12A was added and the plate was wrapped in aluminum foil to prevent illumination. Plates were then pre-incubated in either red or far-red light for 60 minutes before being imaged on an SP8 Leica microscope at 63x magnification. Fluorescent images were gathered every 5 minutes for 30 minutes, using 405 nm excitation with images in two bands (445± 15nm and 525± 25nm). Illumination was switched after 15 minutes to create red to far-red and far-red to red. Generalized polarization (ΔGP) was determined with ImageJ. The intensity at each pixel was compared using the following equation and then normalized to the initial value:

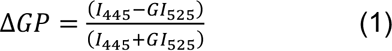

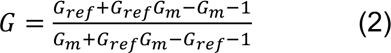

G is the correction factor, G_ref_ is a reference value associated with the dye (0.207), and G_m_ is the measured ΔGP for the dye dissolved in DMSO. The presented protocol was adapted from Owen *et al.* ^52^

### Atomic force microscopy

Cph1-PM-MDA and MDA-MB-231 cells were seeded at 5x10^5^ cells into a Fluorodish (WPI) and incubated overnight with 5 µM PCB and 1800 μg/mL G418 at 37 °C and 5% CO2. A wound was created with a 200 mL tip and washed twice with PBS. The cells were incubated with DMEM with 10% FBS for 1-2 hours. 30min before the AFM experiment, the media was replaced with DMEM containing 2% FBS. Throughout AFM measurements, Cph1-PM-MDA cells were continuously activated or inactivated using 635nm or 740nm light. qp-SCONT cantilevers (Nanosensors) were mounted on a CellHesion 200 AFM (Bruker), connected to an Eclipse Ti inverted light microscope (Nikon). Cantilevers were calibrated using the contact-based approach, followed by coating with 3 mg/ml Concanavalin A (Sigma) for 1 hour at 37 °C. Cantilevers were washed once with PBS before the measurements. Apparent membrane tension was estimated using static tether pulling as follows: Approach velocity was set to 0.5 µm/s, with a contact force of 200 nN, and contact time was varied between 100 ms to 10 s, aiming at maximizing the probability of extruding single tethers. To ensure tether breakage at 0 velocity, the cantilever was then retracted for 10 µm at a velocity of 10 µm/s. Afterwards, the Z-position was kept constant for 25 sec and tether force at the moment of tether breakage was recorded at a sampling rate of 2000 Hz. The resulting force-time curves were analyzed using JPK Data Processing Software. Measurements were run at 37 °C with 5% CO_2_ and samples were used no longer than 2 h for data acquisition. More details can be found here: Bergert *et al.* ^30^

### Phosphatidic acid assay

Total phosphatidic acid was measured with an assay kit purchased from Cell Biolabs, Inc. and run according to the manufacturer’s instructions. In brief, 5x10^6^ cells were plated in triplicate with PCB and either with or without PLD2 modulating drugs, and the plates were then cultured overnight. Plates were illuminated for 4 hours before plates were triple washed for 5 minutes each with ice-cold PBS and then cells were collected via scraping over ice. Cells were pelleted and resuspended in 1 mL of PBS before being sonicated for 30 sec at 1 sec pulses at 30% power. Phosphatidic acid was then removed via phase separation and dried. The dried sample was then resuspended in an assay buffer, and the concentration of phosphatidic acid was determined via fluorescence intensity using excitation at 545±15nm and emission at 590±5nm.

### Antibody staining

5 x10^5^ cells were plated onto 22 x 22 mm glass coverslips placed in each well of a 6-well plated and incubated overnight with PCB at 37 °C and 5% CO_2_. A vertical wound was formed, using a 1 mL pipet tip, the coverslips were twice washed with PBS and incubated with growth media with G418 and PCB 5 µM under the proper illumination condition for 4 hours. And then, they were fixed with 4% paraformaldehyde. Coverslips were then blocked with blocking and PERM buffer (0.2% BSA + 0.1% Triton X-100 in PBS) for 60 minutes at room temperature. Then, the coverslips were incubated with 800:1 rabbit-anti-YAP antibody (#14074, Cell signaling), 200:1 rabbit-anti-pERM antibody (#3726, Cell Signaling), 100:1 mouse-anti-Vinculin monoclonal antibody (13- 9777-82, Invitrogen) in blocking and PERM buffer overnight at 4 °C. Coverslips were triple washed with blocking and PERM buffer and incubated with blocking and PERM buffer containing secondary antibody, 1000:1 Alexa 488 anti-rabbit (#4412, Cell Signaling) or 1000:1 Alexa 488 anti-mouse (#A11029, Invitrogen), 0.1 µg/ml of Phalloidin-TRITC (#ab176757, Abcam), and 1 µg/ml DAPI (#D1306, Invitrogen) for 2 hours at room temperature. The secondary was removed, and the coverslips were triple-washed for 5 minutes each with PBS and mounted on glass slides for imaging.

### Western blot

Cph1-PM-MDA cells were seeded and grown in 150mm Petri dishes (Corning) to 80% confluence with 1800 μg/mL G418 at 37 °C and 5% CO_2_. The cells were then incubated for 24 hours under red or far-red light with 5 µM PCB. Before harvesting, cells were treated with pervanadate (50mM) for 20 minutes. The plates were put on ice and the cells were triple-washed with ice-cold PBS. The cells were then lifted with a rubber policeman and centrifuged for 5 minutes at 600 g. The remaining supernatant was discarded, and cells were frozen overnight to initiate cell lysis. Cells were then defrosted and fully lysed in RIPA buffer containing protease inhibitor cocktail (Sigma). The cells were then sonicated for 20 seconds at 1 sec pulses. Protein concentration was then determined by Bradford assay (Thermo Scientific) according to the manufacturer’s instructions. The protein samples were boiled at 95 °C for 5 minutes, and 40 µg protein samples were loaded and run on 10% Bis-tris SDS-PAGE at 100 V for 90 min. Next, proteins were transferred to nitrocellulose membrane (Carl Roth) at 25 V for 90 min. The membranes were blocked with 5% milk in TBST at room temperature for 60 min and probed with 1:1000 mouse-anti-E-cadherin antibody (#14472, Cell Signaling), 1:1000 rabbit-anti-Vimentin antibody (#5741, Cell Signaling), and 1:2000 rabbit-anti-β-actin antibody (#4970, Cell Signaling) overnight at 4 °C. Primary was removed and the membranes were triple-washed with TBST, and 1:1000 HRP-based anti-mouse secondary antibody (#7076, Cell signaling) and 1:1000 HRP-based anti-rabbit secondary antibodies (#7074, Cell Signaling) were used for chemiluminescent evaluation at room temperature for 2 hours. Pierce ECL western blotting substrate (Thermo Scientific) was used for the developing process.

### Gene expression analysis

Cph1-PM-MDA cells were seeded at 1.6 x10^5^ cells into 45 mm Petri dishes (Corning) and incubated overnight with 5 µM PCB and 1800 μg/mL G418 at 37 °C and 5% CO_2_. Following, Cph1-PM-MDA cells were illuminated for 24 hours at 37 °C and 5% CO_2_. RNA extraction was performed using TRIzol reagent (Life Technologies), following the manufacturer’s protocol. Subsequently, 1 μg of RNA was reverse-transcribed to cDNA with iScript cDNA Synthesis kit (Biorad). Azure Cielo 6 (Azure Biosystem) was used to detect the signal of mRNA expression level with QuiantiNova SYBR Green (Qiagen). Target genes included ZEB1 and FN1 for mesenchymal epithelial transition markers and TEAD1 and CTGF for Yap-associated markers. Primer sequences are provided in **Supplementary Table 1**. GAPDH was used as housekeeping gene. Relative fold changes were calculated by applying the 2^(-ΔΔCt)^ method ^53^.

### Statistics

Statistical analyses were conducted to ensure data normality using the Shapiro-Wilk test. When data followed a normal distribution, a two-sample Student’s t-test (two-tailed) or a pair-sample Student’s t-test (two-tailed) was used for pairwise comparisons; otherwise, a two-sample Mann-Whitney U-test was utilized. For comparisons involving multiple groups, a one-way analysis of variance with Bonferroni’s post-hoc analysis was performed. Statistical analyses were carried out using OriginPro 2020 version 9.7.0.185. The results are presented as mean values with accompanying standard deviations (mean ± SD).

### Data availability

All data in this paper are available. Source data are also provided with this paper.

## Acknowledgements

This work was funded by the European Research Council ERC Starting Grant ARTIST (# 757593, S.V.W) and Deutsche Forschungsgemeinschaft (DFG, German Research Foundation, – Project-ID: 386797833 – SFB 1348: Dynamic Cellular Interphases and GA2268/4-1).

## Author contributions

B.M.B., J.P. and S.V.W. designed the experiments. B.M.B. and J.P. performed the experiments, analyzed the data and generated the figures. M.B. and A.D. supported the atomic force microscope measurements. C.T. and M.G. conducted the analysis of single-cell tracks. All authors discussed data and wrote the manuscript.

## Supplementary Information

### SUPPLEMENTARY MOVIES

**Supplementary Movie 1:** Cellular migration for wound-healing assay of MDA and MCF-7 cells. Frame rate (20 fps, with 20 frames per second). Scale-bar: 100 μm.

**Supplementary Movie 2:** Cellular migration for wound-healing assay of Cph1-PM-MDA cells under far-red and red light. Frame rate (20 fps, with 20 frames per second). Scale-bar: 100 μm.

**Supplementary Movie 3:** Single-cell tracking of individual cell nuclei of Cph1-PM-MDA cells under far-red or red light. Frame rate (20 fps, with 20 frames per second). Scale-bar: 100 μm.

**Supplementary Movie 4:** Cellular migration for wound-healing assay of Cph1-PM-MDA cells under far-red or red light in the presence of VU. Frame rate (20 fps, with 20 frames per second). Scale-bar: 100 μm.

**Supplementary Movie 5:** Cellular migration for wound-healing assay of Cph1-PM-MDA cells under far-red or red light in the presence of Oleic acid. Frame rate (20 fps, with 20 frames per second). Scale-bar: 100 μm.

---

**Supplementary Figure 1:**
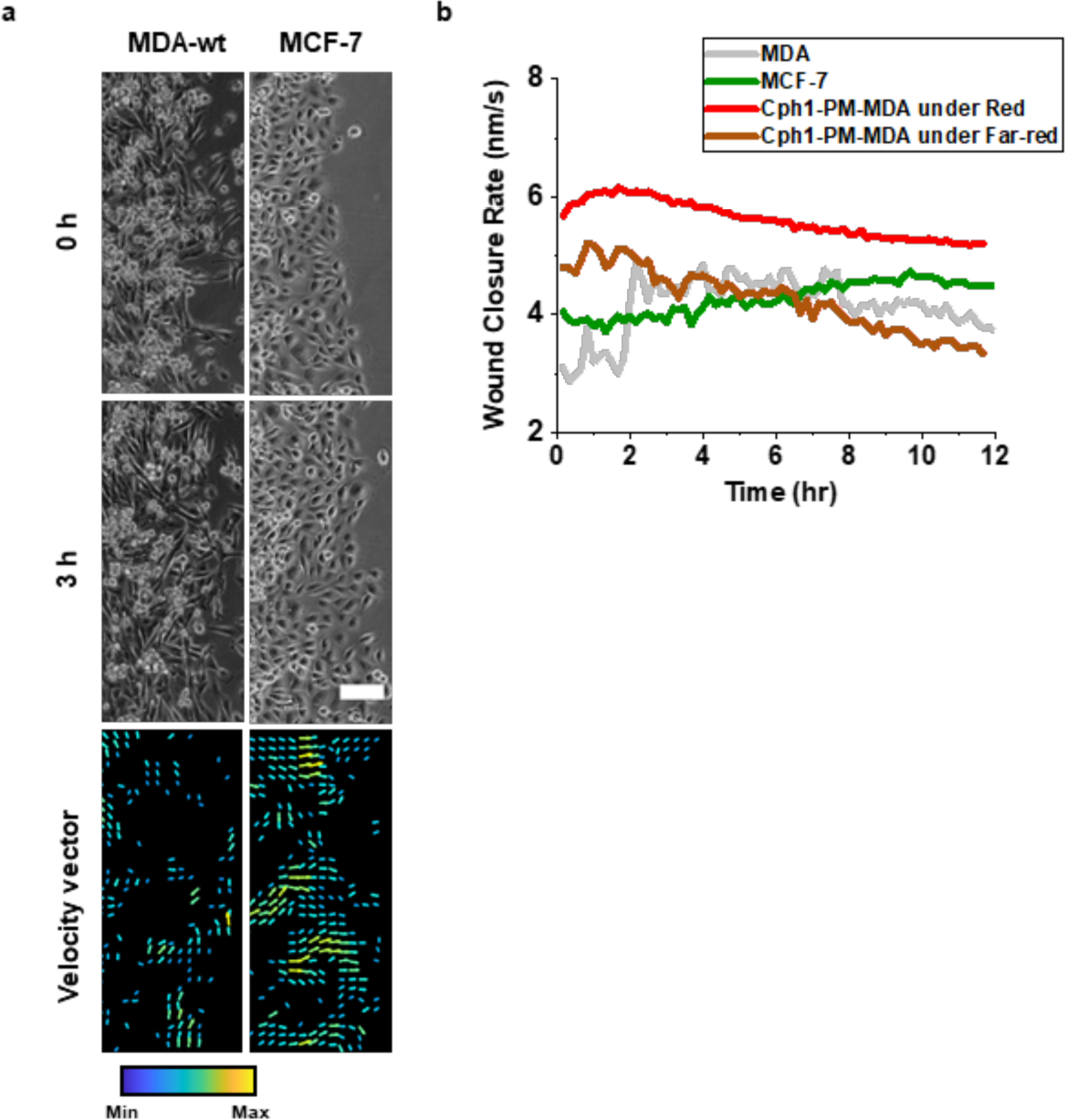
Migration data for individual and collective migration. (a) Wound healing migration patterns for MDA-MB-231 (MDA) and MCF-7. **(b)** Velocity histograms depicting cellular migration patterns for MDA and MCF-7. Scale bar (a), 100 μm.

**Supplementary Figure 2:**
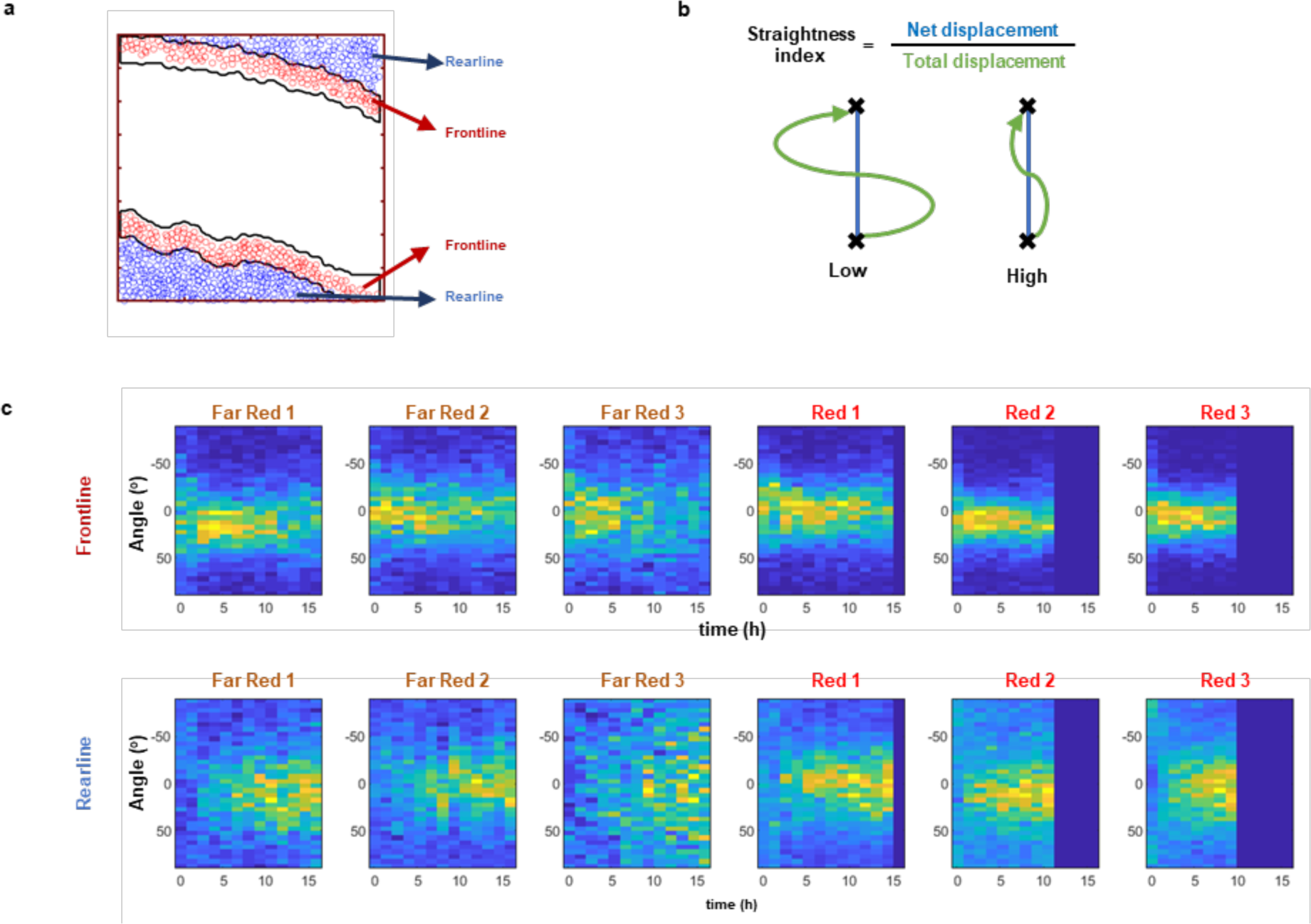
Analysis of single-cell tracking data. (a) The regions were divided into frontline of around 3 lines and rearline of rest region. **(b)** The schematic of straightness index was presented. Straightness index values are calculated by dividing net displacement into total displacement. **(c)** The plots show the migration angle of individual Cph1-PM-MDA cells in the frontlines and rearlines under red and far- red light for 16 hours. The number indicates the technical replications.

**Supplementary Figure 3:**
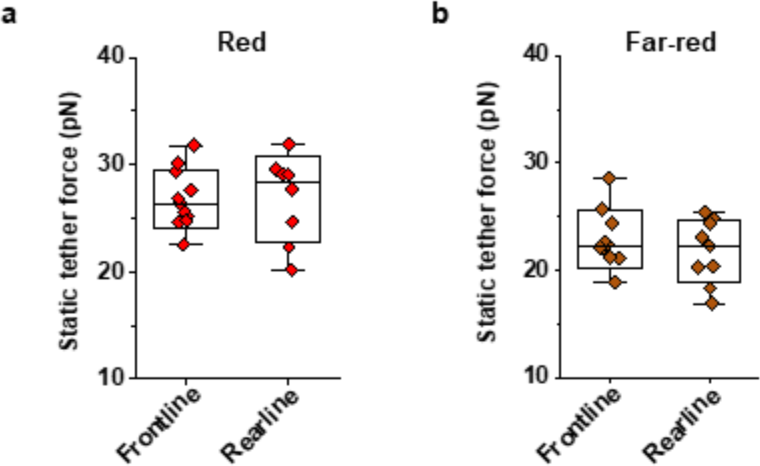
Static tether force of Cph1-PM-MDA cells. Static tether forces were measured after exposures to either red or far-red light tension in the frontline and rearline in the wound direction both under red (a) and far-red light (b). Data source and statistic details are provided as a source data file.

**Supplementary Figure 4:**
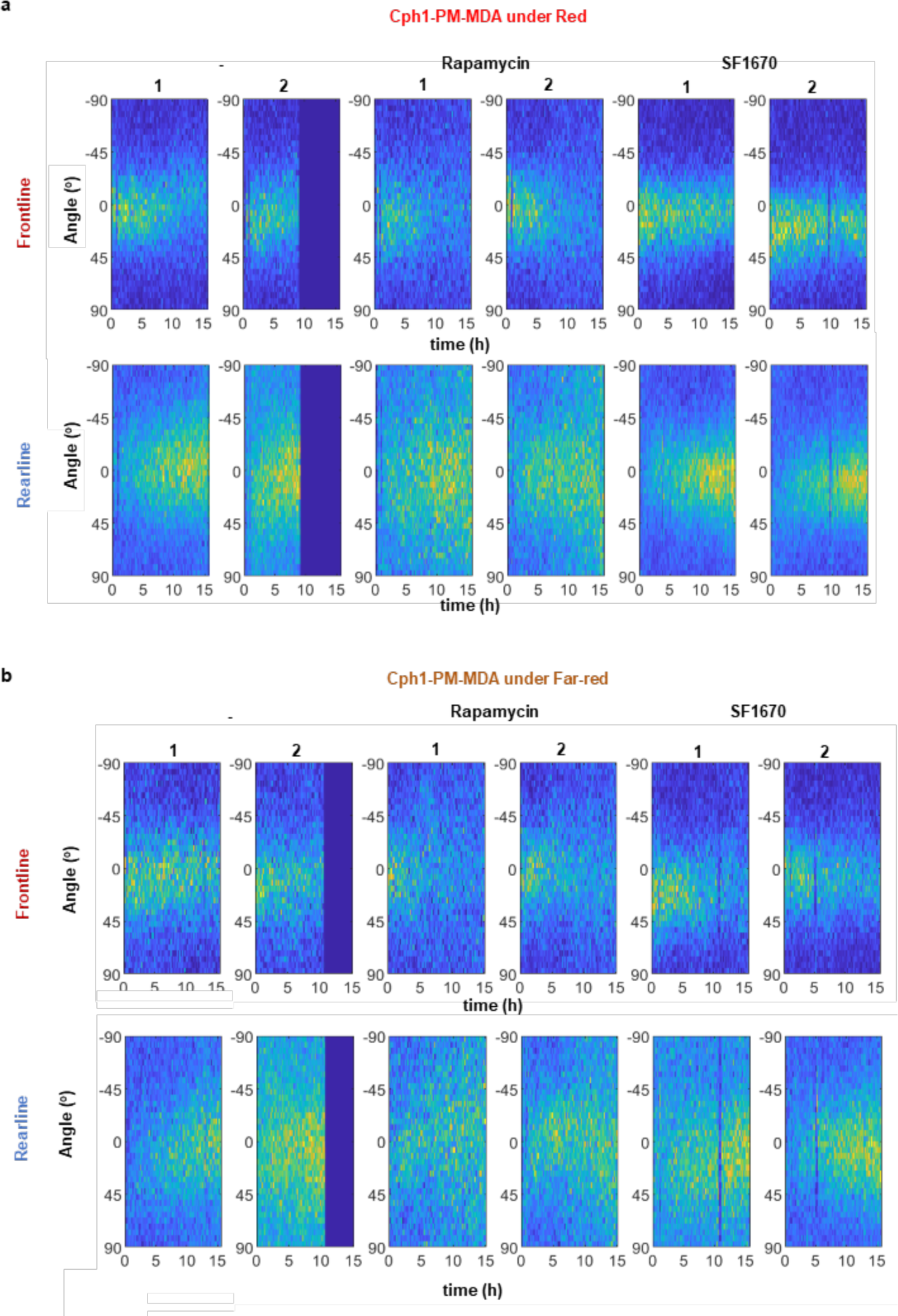
Migration angles of Cph1-PM-MDA cells without or with rapamycin and SF1670. (a, b) The plots show the migration angle of individual Cph1-PM-MDA cells in the frontlines and rearlines under red and far-red light for 16 hours without treat or with treatment of rapamycin or SF1670. The number indicate the technical replications.

**Supplementary Table 1:**
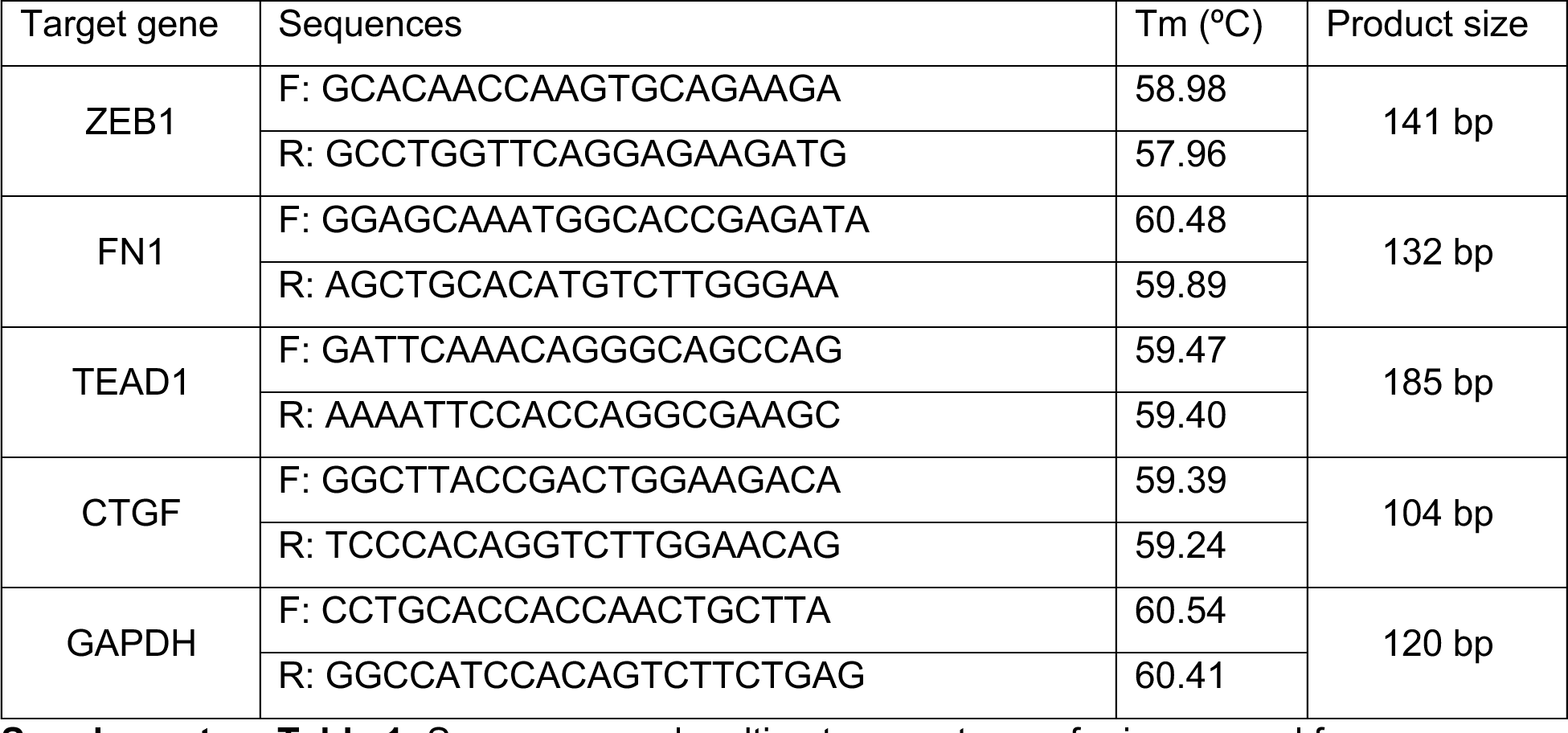
Sequences and melting temperatures of primers used for gene expression analysis.

## References

1. Yamada KM, Sixt M. Mechanisms of 3D cell migration. Nature Reviews molecular cell biology 20, 738–752 (2019).

2. Canty L, Zarour E, Kashkooli L, François P, Fagotto F. Sorting at embryonic boundaries requires high heterotypic interfacial tension. Nature communications 8, 157 (2017).

3. Cockburn K, Rossant J. Making the blastocyst: lessons from the mouse. The Journal of clinical investigation 120, 995–1003 (2010).

4. Friedl P, Locker J, Sahai E, Segall JE. Classifying collective cancer cell invasion. Nature cell biology 14, 777–783 (2012).

5. Scarpa E, Mayor R. Collective cell migration in development. Journal of Cell Biology 212, 143–155 (2016).

6. Friedl P, Mayor R. Tuning collective cell migration by cell–cell junction regulation. Cold Spring Harbor perspectives in biology 9, a029199 (2017).

7. Leckband DE, de Rooij J. Cadherin adhesion and mechanotransduction. Annual review of cell and developmental biology 30, 291–315 (2014).

8. Hayer A, et al. Engulfed cadherin fingers are polarized junctional structures between collectively migrating endothelial cells. Nature cell biology 18, 1311–1323 (2016).

9. Nzigou Mombo B, et al. Reversible photoregulation of cell-cell adhesions with opto-E-cadherin. Nature Communications 14, 6292 (2023).

10. Bertocchi C, et al. Nanoscale architecture of cadherin-based cell adhesions. Nature cell biology 19, 28–37 (2017).

11. Tepass U, Truong K, Godt D, Ikura M, Peifer M. Cadherins in embryonic and neural morphogenesis. Nature reviews Molecular cell biology 1, 91–100 (2000).

12. Takeichi M. Cadherin cell adhesion receptors as a morphogenetic regulator. Science 251, 1451–1455 (1991).

13. Nagafuchi A, Takeichi M. Cell binding function of E-cadherin is regulated by the cytoplasmic domain. The EMBO Journal 7, 3679–3684 (1988).

14. Bianchini JM, Kitt KN, Gloerich M, Pokutta S, Weis WI, Nelson WJ. Reevaluating αE-catenin monomer and homodimer functions by characterizing E-cadherin/αE-catenin chimeras. Journal of Cell Biology 210, 1065–1074 (2015).

15. Raucher D, Sheetz MP. Cell spreading and lamellipodial extension rate is regulated by membrane tension. The Journal of cell biology 148, 127–136 (2000).

16. Gurtovenko AA, Anwar J. Modulating the structure and properties of cell membranes: the molecular mechanism of action of dimethyl sulfoxide. J Phys Chem B 111, 10453–10460 (2007).

17. Yumura S, Fukui Y. Filopodelike projections induced with dimethyl sulfoxide and their relevance to cellular polarity in Dictyostelium. The Journal of cell biology 96, 857–865 (1983).

18. Braig S, Schmidt, S., Stoiber, K., Händel, C., Möhn, T., Werz, O., Müller, R., Zahler, S., Koeberle, A., Käs, J.A. and Vollmar, A.M. Pharmacological targeting of membrane rigidity: implications on cancer cell migration and invasion. New Journal of Physics 17, (2015).

19. Sitarska E, Diz-Muñoz A. Pay attention to membrane tension: Mechanobiology of the cell surface. Current opinion in cell biology 66, 11–18 (2020).

20. Diz-Muñoz A, et al. Membrane tension acts through PLD2 and mTORC2 to limit actin network assembly during neutrophil migration. PLoS biology 14, e1002474 (2016).

21. Loh J, et al. An acute decrease in plasma membrane tension induces macropinocytosis via PLD2 activation. Journal of Cell Science 132, jcs232579 (2019).

22. Egea-Jimenez AL, Zimmermann P. Phospholipase D and phosphatidic acid in the biogenesis and cargo loading of extracellular vesicles. J Lipid Res 59, 1554–1560 (2018).

23. Disanza A, et al. CDC42 switches IRSp53 from inhibition of actin growth to elongation by clustering of VASP. The EMBO journal 32, 2735–2750 (2013).

24. Echarri A, et al. An Abl-FBP17 mechanosensing system couples local plasma membrane curvature and stress fiber remodeling during mechanoadaptation. Nature communications 10, 5828 (2019).

25. Rasoulinejad S, Mueller M, Nzigou Mombo B, Wegner SV. Orthogonal blue and red light controlled cell–cell adhesions enable sorting-out in multicellular structures. ACS Synthetic Biology 9, 2076–2086 (2020).

26. Vitorino P, Meyer T. Modular control of endothelial sheet migration. Genes & development 22, 3268–3281 (2008).

27. Nieman MT, Prudoff RS, Johnson KR, Wheelock MJ. N-cadherin promotes motility in human breast cancer cells regardless of their E-cadherin expression. The Journal of cell biology 147, 631–644 (1999).

28. Mbalaviele G, Dunstan CR, Sasaki A, Williams PJ, Mundy GR, Yoneda T. E-cadherin expression in human breast cancer cells suppresses the development of osteolytic bone metastases in an experimental metastasis model. Cancer research 56, 4063–4070 (1996).

29. Zhang Y-L, Frangos JA, Chachisvilis M. Laurdan fluorescence senses mechanical strain in the lipid bilayer membrane. Biochemical and biophysical research communications 347, 838–841 (2006).

30. Bergert M, Diz-Muñoz A. Quantification of Apparent Membrane Tension and Membrane-to- Cortex Attachment in Animal Cells Using Atomic Force Microscopy-Based Force Spectroscopy. In: Mechanobiology: Methods and Protocols (ed Zaidel-Bar R). Springer US (2023).

31. Liu Y, et al. Constitutively active ezrin increases membrane tension, slows migration, and impedes endothelial transmigration of lymphocytes in vivo in mice. *Blood*, The Journal of the American Society of Hematology 119, 445–453 (2012).

32. Tsujita K, et al. Homeostatic membrane tension constrains cancer cell dissemination by counteracting BAR protein assembly. Nature Communications 12, 5930 (2021).

33. Pontes B, et al. Membrane tension controls adhesion positioning at the leading edge of cells. Journal of Cell Biology 216, 2959–2977 (2017).

34. Carisey A, et al. Vinculin regulates the recruitment and release of core focal adhesion proteins in a force-dependent manner. Current biology 23, 271–281 (2013).

35. Case LB, et al. Molecular mechanism of vinculin activation and nanoscale spatial organization in focal adhesions. Nature Cell Biology 17, 880–892 (2015).

36. Kim D-H, Wirtz D. Focal adhesion size uniquely predicts cell migration. Biophysical Journal 104, 319a (2013).

37. Benham-Pyle BW, Pruitt BL, Nelson WJ. Mechanical strain induces E-cadherin–dependent Yap1 and β-catenin activation to drive cell cycle entry. Science 348, 1024–1027 (2015).

38. Fang Y, Vilella-Bach M, Bachmann R, Flanigan A, Chen J. Phosphatidic acid-mediated mitogenic activation of mTOR signaling. Science 294, 1942–1945 (2001).

39. Sozzani S, et al. Propranolol, a phosphatidate phosphohydrolase inhibitor, also inhibits protein kinase C. Journal of Biological Chemistry 267, 20481–20488 (1992).

40. Lavieri R, et al. Design and synthesis of isoform-selective phospholipase D (PLD) inhibitors. Part II. Identification of the 1, 3, 8-triazaspiro [4, 5] decan-4-one privileged structure that engenders PLD2 selectivity. Bioorganic & medicinal chemistry letters 19, 2240-2243 (2009).

41. Sens P, Plastino J. Membrane tension and cytoskeleton organization in cell motility. Journal of Physics: Condensed Matter 27, 273103 (2015).

42. Żelasko J, Czogalla A. Selectivity of mTOR-Phosphatidic Acid Interactions Is Driven by Acyl Chain Structure and Cholesterol. Cells 11, 119 (2021).

43. Yano H, et al. Inhibition of histamine secretion by wortmannin through the blockade of phosphatidylinositol 3-kinase in RBL-2H3 cells. Journal of Biological Chemistry 268, 25846–25856 (1993).

44. Li Y, et al. Pretreatment with phosphatase and tensin homolog deleted on chromosome 10 (PTEN) inhibitor SF1670 augments the efficacy of granulocyte transfusion in a clinically relevant mouse model. *Blood*, The Journal of the American Society of Hematology 117, 6702–6713 (2011).

45. Sarbassov DD, et al. Prolonged rapamycin treatment inhibits mTORC2 assembly and Akt/PKB. Molecular cell 22, 159–168 (2006).

46. Peglion F, et al. PTEN inhibits AMPK to control collective migration. Nature Communications 13, 4528 (2022).

47. Rübsam M, Broussard JA, Wickström SA, Nekrasova O, Green KJ, Niessen CM. Adherens junctions and desmosomes coordinate mechanics and signaling to orchestrate tissue morphogenesis and function: an evolutionary perspective. Cold Spring Harbor perspectives in biology 10, a029207 (2018).

48. Huber M, Casares-Arias J, Fässler R, Müller DJ, Strohmeyer N. In mitosis integrins reduce adhesion to extracellular matrix and strengthen adhesion to adjacent cells. Nature Communications 14, 2143 (2023).

49. Buckley CD, et al. The minimal cadherin-catenin complex binds to actin filaments under force. Science 346, 1254211 (2014).

50. Del Rio A, Perez-Jimenez R, Liu R, Roca-Cusachs P, Fernandez JM, Sheetz MP. Stretching single talin rod molecules activates vinculin binding. Science 323, 638–641 (2009).

51. Price AJ, Cost A-L, Ungewiß H, Waschke J, Dunn AR, Grashoff C. Mechanical loading of desmosomes depends on the magnitude and orientation of external stress. Nature communications 9, 5284 (2018).

52. Owen DM, Rentero C, Magenau A, Abu-Siniyeh A, Gaus K. Quantitative imaging of membrane lipid order in cells and organisms. Nature Protocols 7, 24–35 (2012).

53. Livak KJ, Schmittgen TD. Analysis of relative gene expression data using real-time quantitative PCR and the 2− ΔΔCT method. methods 25, 402–408 (2001).

